# H-MAGMA, inheriting a shaky statistical foundation, yields excess false positives

**DOI:** 10.1101/2020.08.20.260224

**Authors:** Ronald Yurko, Kathryn Roeder, Bernie Devlin, Max G’Sell

## Abstract

The ‘snp-wise mean model’ of Multi-marker Analysis of GenoMic Annotation is often used to perform gene-level testing for association with disease and other phenotypes. This methodology, in turn, forms the foundation for H-MAGMA. Unfortunately, that foundation is unsound, with implications for H-MAGMA results published in *Nature Neuroscience* regarding genes associated with psychiatric disorders: e.g., only 125 of H-MAGMA’s 275 reported discoveries for autism replicate when the foundation’s flaws are corrected.

---

The ‘snp-wise mean model’ of Multi-marker Analysis of GenoMic Annotation^1^ (hereafter MAGMA) is often used to perform gene-level testing for association with disease or other phenotypes, taking as input genomewide association study (GWAS) summary statistics and reference linkage disequilibrium (LD) data. The success of this methodology (MAGMA has 826 Google Scholar citations as of 18 August, 2020) has led to its incorporation into a variety of tools, including an atlas of over 4,700 GWAS results^2^ and various testing methods. For example, it is the foundation for H-MAGMA^3^, which also incorporates Hi-C data and uses the Benjamini-Hochberg (BH)^4^ procedure for false discovery rate (FDR) control. When applying MAGMA, however, we noted that its distributional properties did not comport with statistical expectation.

When investigating the basis of MAGMA’s distributional properties, we also discovered a critical departure of the accepted MAGMA implementation from the manuscript details. MAGMA, as described in the manuscript, builds on Brown’s approximation of Fisher’s method for combining dependent SNP-level p-values^5^, adjusting for the LD-induced covariance of SNP p-values. This statistical approximation, however, is *valid only for one-sided tests*. GWAS summary statistics are necessarily two-sided tests because which SNP allele confers risk is not known, a priori^6^. When applied to two-sided tests, as in the analysis of GWAS summary statistics by MAGMA, the assumed null distribution is incorrect in both its distributional form and its covariance. The correct null distribution for simulated multivariate normal two-sided test statistics (Fig. 1a) does not follow the re-scaled chi-square distribution implied by Brown’s approximation (denoted by *MAGMA: paper)*. Furthermore, comparison to a known correction to the covariance for two-sided tests^7^ leads to a stark difference with the one-sided test approximation (Supplementary Fig. 1; see Supplement for details of this and other calculations).

**Fig. 1.**
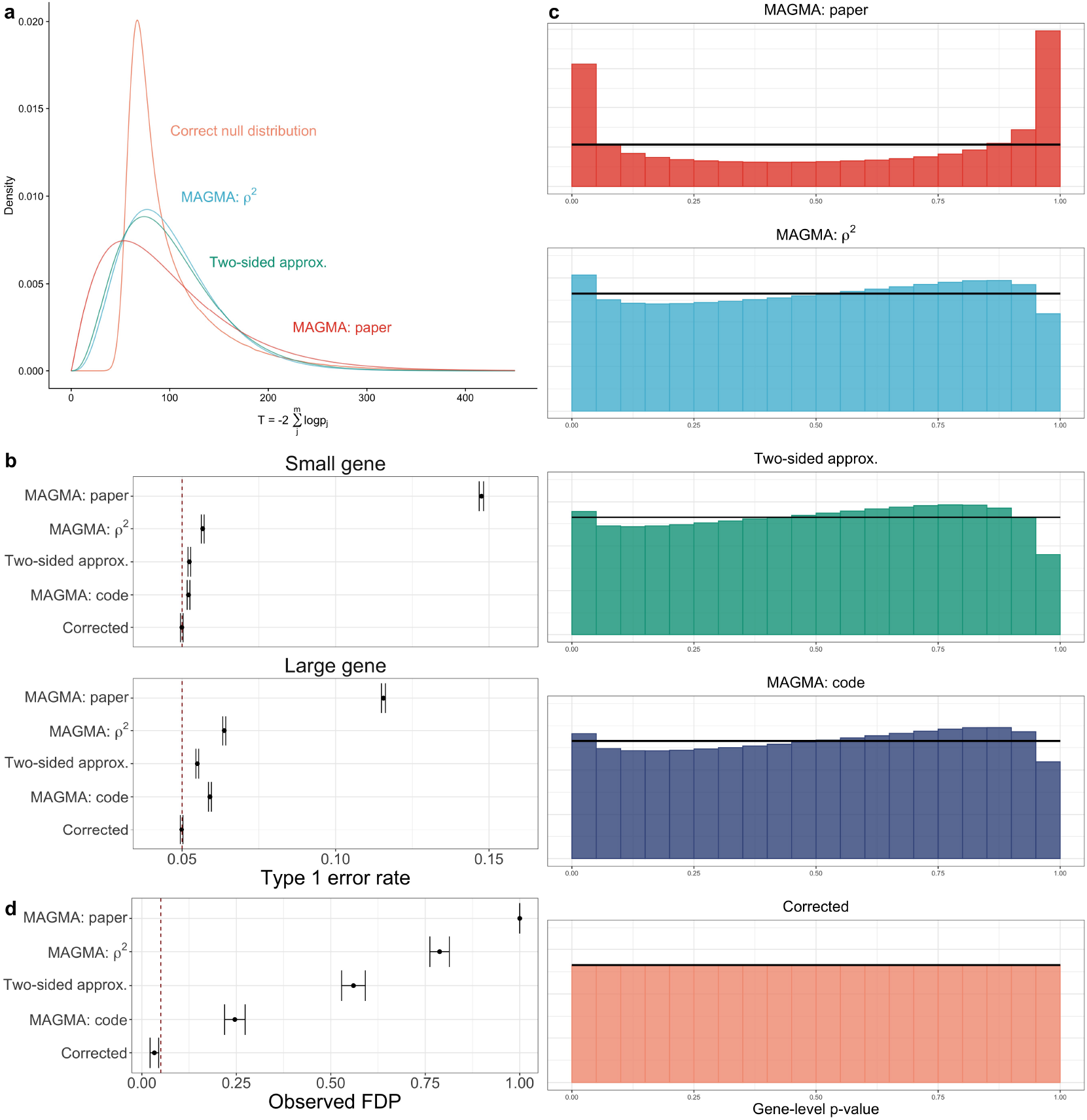
**a**, Comparison of the different covariance approximations for fitting the test statistic distribution with Brown’s rescaled *χ*^2^ using two-sided summary statistics. Test statistic distribution is generated using fifty-dimensional multivariate Gaussian distribution centered with mean zero and block correlation *ρ* = 0.50. **b**, Comparison of the type 1 error rate control, plus/minus two standard errors at target *α* = 0.05 (denoted by dashed red line) by gene-level method for small (17 SNPs) and large (1,068 SNPs) example genes. **c**, Comparison of null histograms by method for ≈ 53000 genes of different sizes averaged over 1,000 simulations (standard errors are too small to be visible). Horizontal black lines represents ideal uniform distribution for null p-values. **d**, Comparison of average BH false discovery proportions (FDP) across 1000 simulations, plus/minus two standard errors, at target FDR level *α* = 0.05 (denoted with dashed red line) by gene-level method.

This incorrect null distribution should lead to invalid and non-uniform null p-values with a severely inflated error rate. Curiously, this does not comport with the observed performance of MAGMA when tests are conducted at small significance level *α*. How could this be? The software embodies two undocumented, ad-hoc corrections: replacing the correlation coefficient *ρ* in Brown’s approximation with its square, which acts as a rough correction for one-versus two-sidedness (denoted as *MAGMA: ρ*^2^ in Fig. 1), followed by an empirically-motivated warping of the p-values to reduce the false positive rate (*MAGMA: code*). While these corrections together result in improved error rate control at small *α*, ultimately, this extension of Brown’s approximation is invalid for two-sided tests (Fig. 1a) and it yields an inflated error rate that worsens for larger genes (Fig. 1b and Supplementary Fig. 2).

MAGMA’s incorrect null p-value distributions (Fig. 1c) are particularly relevant for procedures that correct for multiple testing, such as family-wise error rates (Supplementary Fig. 3 and 4). We demonstrate this impact with simulations using real genotype data^8^ to show its failure to maintain FDR control using the BH procedure (Fig. 1d). Additionally, we observe that the concern for MAGMA in the setting of gene-set enrichment analysis is a loss of power due to the properties of the procedures (see *Supplement* and Supplementary Fig. 5–7). In comparison, Monte Carlo-based approaches to computing the null guarantee appropriate error-rate control and uniform *p*-value distributions under the assumed model (Fig. 1b-d), even when using the same test statistic as MAGMA (referred to as *Corrected*).

The use of MAGMA impacts the results and development of new procedures that inherit its statistical flaws. We examine the results of one such paper, published in *Nature Neuroscience*, which proposes H-MAGMA^3^. After introducing the concept of H-MAGMA, which relies on MAGMA for computing gene-level p-values and BH for FDR control, the authors apply it to data from five psychiatric disorders. Reanalyzing these data, we observe that the H-MAGMA p-value distributions are improper (Fig. 2a) and testing using corrected p-values yields substantially smaller subsets of the reported results (Fig. 2b and Supplementary Table 1). This reduction is more prevalent in the weaker signal setting for autism spectrum disorder, for which only 125 of the 275 genes H-MAGMA associates with autism replicate with the corrected p-values.

**Fig. 2.**
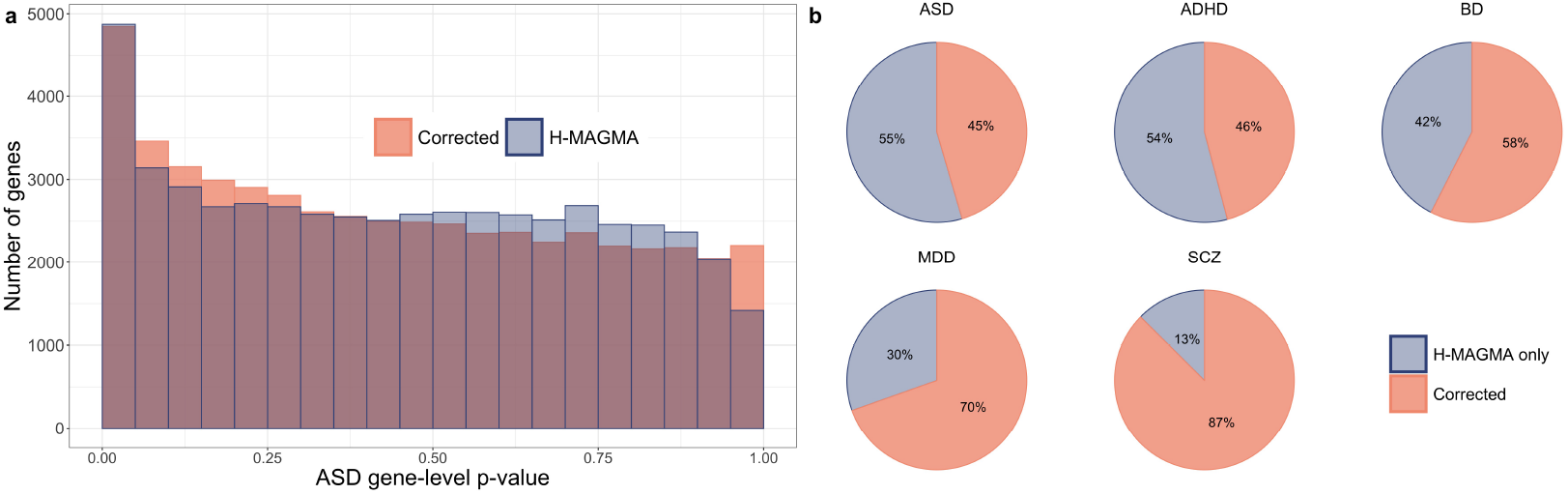
**a**, Comparison of autism spectrum disorder (ASD) gene-level p-values, based on the adult Hi-C data annotations, computed with H-MAGMA versus corrected Monte Carlo-based approach. **b**, Percentage of H-MAMGA reported discoveries using BH at target FDR level *α* = 0.05 for five psychiatric disorders due to MAGMA inflation versus the remaining Monte Carlo-corrected set.

Correcting MAGMA’s underlying gene-level p-value computation is essential to ensure that novel extensions, such as H-MAGMA, do not yield excess false positives. As our results suggest, a simple solution is to replace Brown’s approximation in MAGMA with Monte Carlo-based procedures, similar to *VEGAS*^9,10^. We believe these results are critical for researchers wishing to interpret gene-based testing or for those wishing to build new methods in this challenging area.

## Data availability

All data used in the manuscript are available at https://github.com/ryurko/HMAGMA-comment.

## Code availability

All code used in the manuscript are available at https://github.com/ryurko/HMAGMA-comment.

## Supplement

### Background

#### Combining p-values under dependence

Fisher’s method^11^ is a classical approach to combine p-values from *m* tests. Given observed p-values *p_j_* for each test *j* ∈ [*m*], it computes a summary test statistic,

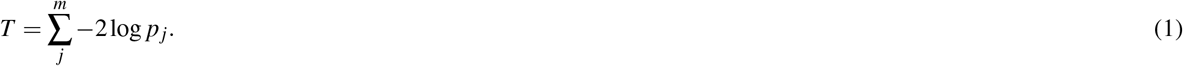

Under the assumption of independent p-values, Fisher’s test statistic 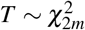. Fisher’s method is used to test the global null, i.e. all SNPs *j* ∈ [*m*] in gene *g* are null versus at least one SNP in the gene is non-null,

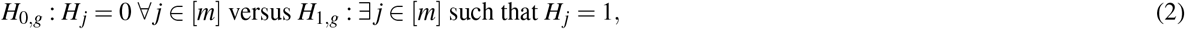

where *H_j_* = 0 if SNP *j* is null, and *H_j_* = 1 if non-null.

However, p-values for nearby SNPs are often dependent due to LD. Brown introduced an extension of Fisher’s method in the case of dependence^5^, for the setting where *m* tests are based on multivariate Gaussian random variables,

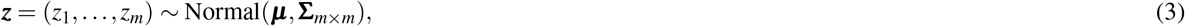

where ***μ*** is the vector of true centers for the *m* tests and **Σ**_*m*×*m*_ is the corresponding covariance matrix. Brown’s test uses the same Fisher’s test statistics as in Equation 1, but assumes that the null distribution is 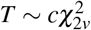, a re-scaled *χ*^2^ distribution. The two constants *c* and *v* are calculated by matching the first two moments of the re-scaled 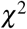 to the moments of Fisher’s test statistic *T* induced by the multivariate Gaussian,

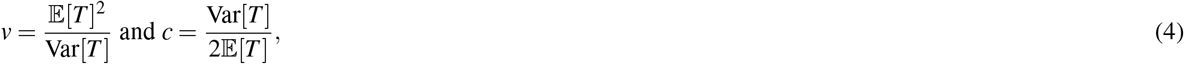

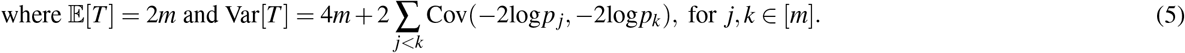

Instead of computing the covariance directly with numerical integration, which can be computationally intensive in-omics settings such as GWAS, Brown approximated the covariance between tests *j* and *k* as a function of the correlation of the corresponding Gaussians, *ρ_jk_*,

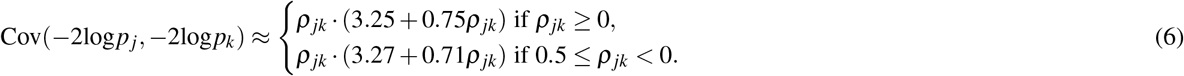

Since Brown’s initial publication, there have been further refinements to this approximation^12,13^; for example, Kost and McDermott^12^ used polynomial regression over a grid of values for −0.98 ≤ *ρ_jk_* ≤ 0.98 by increments of 0.02,

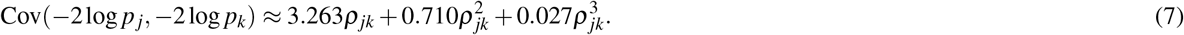

**Supplementary Fig. 1.**
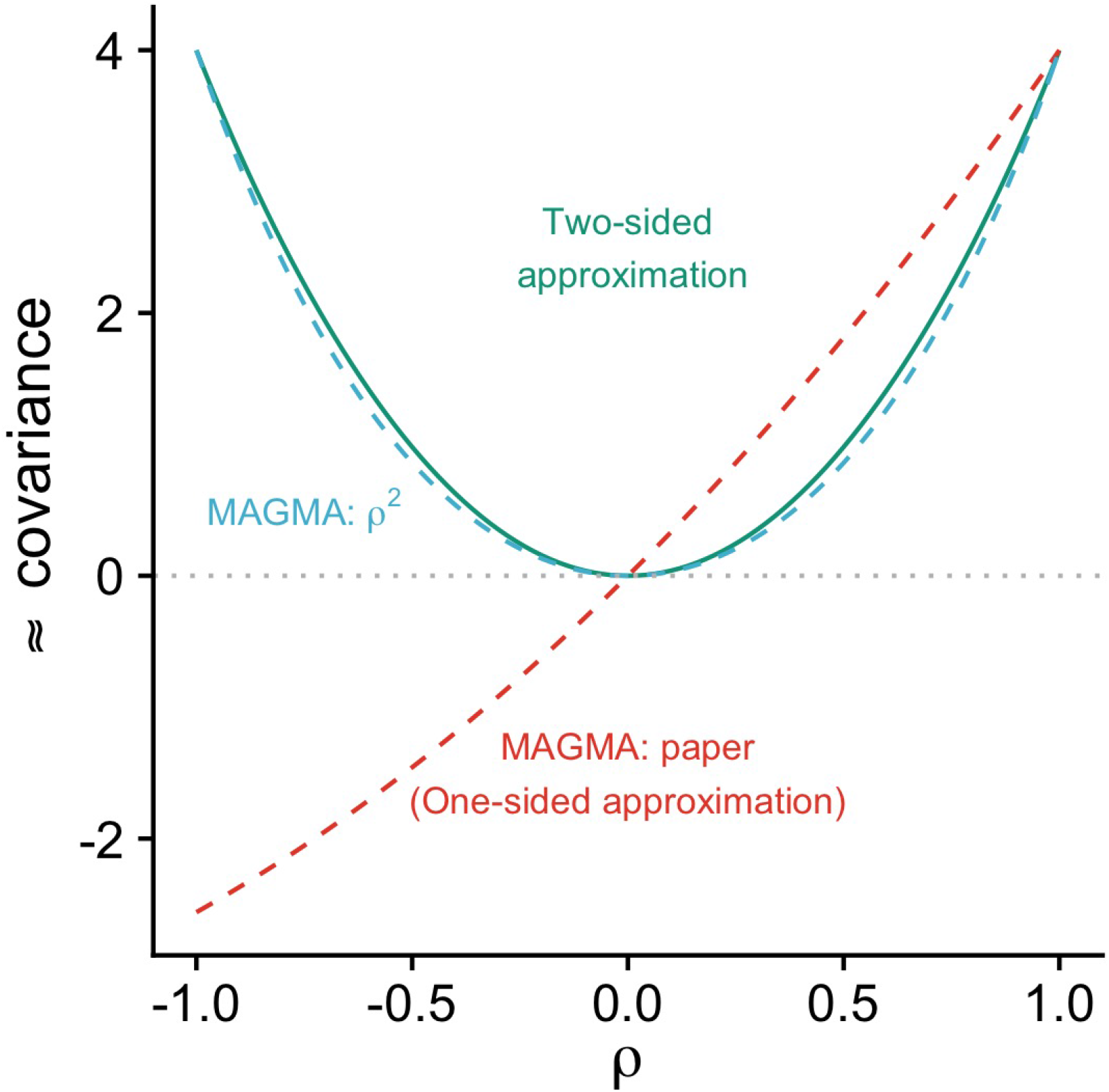
Comparison of covariance approximations for two-sided tests (green-solid) versus Brown’s one-sided approximation described in MAGMA paper (red-dashed) and the code implemented in MAGMA with *ρ*^2^ (cyan-dashed).

However, Brown’s covariance approximation and its various refinements are based on the usage of one-sided tests *p_j_* = 1 − *ϕ*(*z_j_*), where *z_j_* is the corresponding z-statistic for test *j* and *ϕ* is the Gaussian cumulative distribution function. Yang et al.7 introduced an approximation for two-sided tests, *p_j_* = 2*ϕ* (−|*z_j_*|), in the context of detecting association between a SNP and multivariate phenotypic traits, using a tenth order polynomial,

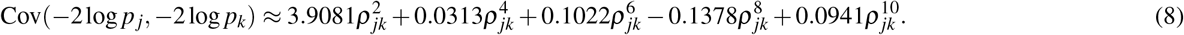

This yields a drastically different covariance approximation from one-sided tests as displayed in Supplementary Fig. 1. We provide a simpler two-sided approximation using polynomial regression over a grid of values for −1 ≤ *ρ_jk_* ≤ 1 by increments of 0.01,

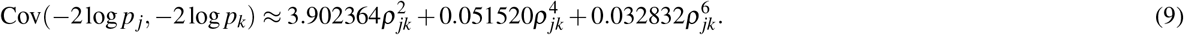

The median difference between the approximations in Equations 8 and 9 is 0.0006465, with a maximum observed difference of 0.011184. Equation 9 also displays a smaller maximum absolute difference of 0.0009217044 with a recent calculation using Hermite polynomials^14^ than Equation 8, which has a maximum absolute difference of 0.0104.

#### MAGMA ‘snp-wise-mean model’

The MAGMA ‘snp-wise-mean model’ is used to compute a gene-level test statistic from GWAS summary statistics. In the original publication^1^, the authors describe the use of Brown’s covariance approximation, which is inappropriate for two-sided tests (Supplementary Fig. 1). However, the maintainers of the MAGMA software have made several adjustments to this approximation not described in the manuscript. First, prior to approximating the covariance based on the sign of the correlation, they square the correlation values, 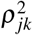, resulting in,

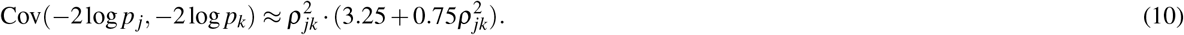

This covariance approximation (Supplementary Fig. 1) alleviates the initial stark difference in covariance values in the presence of negative correlation from using Brown’s one-sided approximation and is much closer to, but still under-estimating, the appropriate approximation for two-sided tests. The software includes an additional adjustment to the resulting gene-level p-value *p_g_*,

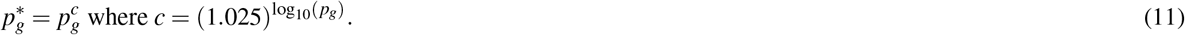

Based on correspondence with the maintainers of MAGMA, they determined the use of an adjusted p-value 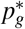 from the power *c* after viewing simulations to reveal Brown’s approximation would yield smaller p-values than the truth. By using the power adjustment above, smaller values *p_g_* will receive a *stronger* adjustment. It is unfortunate that these adjustments were not presented in the manuscript, which thus misleads readers who are exploring the usage of similar methodology for combining p-values from GWAS summary statistics.

### Methodology

#### Methods for computing gene-level p-values

To evaluate the impact of MAGMA’s implementation, we compare these approaches:

1. *MAGMA: paper* - the approach presented in the original MAGMA manuscript following Brown’s covariance approximation for one-sided tests, red-dashed line in Supplementary Fig. 1,
2. *MAGMA: ρ*^2^ - replacement of Brown’s covariance approximation with squared correlation values, cyan-dashed line in Supplementary Fig. 1,
3. *MAGMA: code* - includes the use of both *ρ*^2^ and the adjustment power *c*,
4. *Two-sided approximation* - replacement of Brown’s covariance approximation with the appropriate two-sided covariance^7^, green-solid line in Supplementary Fig. 1.
5. *Corrected* - Monte Carlo simulation using the Fisher test statistic, 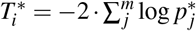. Details below.

The *Corrected* Monte Carlo-based approach for computing empirical p-values differs from MAGMA in that it does not rely on Brown’s original assumption of a re-scaled *χ*^2^ distribution. Instead, we generate *N* draws of *m*-dimensional Gaussian random variables,

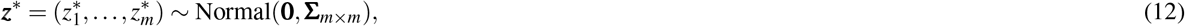

where **0** is a vector of zeroes representing the null distribution with **Σ**_*m*×*m*_ as the gene’s covariance matrix. A test statistic 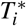 is calculated for each of the *i* ∈ [*N*] draws and, using the observed test statistic *T*, an empirical gene-level p-value is calculated as:

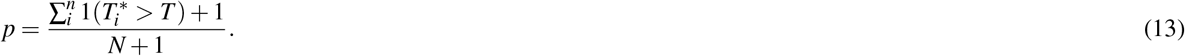

To be consistent with MAGMA, we use the same test statistic 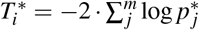, where 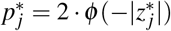 is a two-sided p-value. While the *Corrected* results in this manuscript refers matching the MAGMA test statistic, we observed equivalent results when using *VEGAS*^9,10^, a similar Monte Carlo-based approach based on a different test statistic, 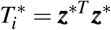.

#### Multivariate Gaussian simulation

As a ‘toy’ example for demonstrating the flaw of Brown’s approximation for two-sided test statistics, we generate null multivariate Gaussian random variables as in Equation 12, which matches the initial assumption made by Brown in Equation 3. For simplicity, we a *m =* 50-dimensional covariance matrix with block correlation *ρ* = 0.5, where the diagonal elements **Σ**_*j,j*_ = 1 and all off-diagonal elements **Σ**_*j,k*_ = 0.5 for *j* ≠ *k*, and *j, k* = 1,…, *m*. We use 1,000,000 simulations under this model to generate the correct null distribution for the test statistic 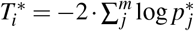 in Fig 1a, where 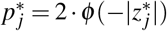 is a two-sided p-value, as compared to Brown’s re-scaled *χ*^2^ distribution with three different covariance approximations: *MAGMA: paper* (red), *MAGMA: ρ*^2^ (cyan), and the *Two-sided approximation* (green).

#### Example genotype data

To evaluate the performance of the gene-level methods with real data, we randomly select genes using the sample of 503 individuals from the 1000 Genomes project^8^ of European ancestry (build 37), with NCBI gene-locations accessed from the MAGMA site: https://ctg.cncr.nl/software/magma. After standard pre-processing steps, excluding SNPs with minor allele frequency (MAF) less than 0.05 and call rate less than 0.95, the remaining SNPs were assigned to 19,326 genes using the MAGMA ‘annotate’ command with 10kb padding (upstream and downstream). We randomly select fifteen candidate genes from each autosomal chromosome, with stratified sampling of five genes from each of the following size groups: (1) [10,100], (2) (100,500], and (3) (500, ∞) SNPs. To remove SNPs displaying high LD values, these candidate genes were pruned in the following manner:

1. order the SNPs in the gene by minor allele frequencies (MAFs) in descending order,
2. starting with the SNP with the highest MAF,

- remove all SNPs with *r*^2^ ≥ 0.95 within the gene,
- move on to the next SNP that is still remaining,
3. return the retained SNPs that are not in high LD for simulations.

We then randomly selected nine example genes, with stratified sampling of three genes from the same size groups as above: (1) [10,100], (2) (100,500], and (3) (500, ∞) SNPs remaining after pruning. The nine randomly selected genes (with *m* SNPs after pruning) are: KRT1 (*m* = 17), FAM13B (24), BMPR2 (38), IDO2 (101), PREX2 (218), RAI14 (220), RYR3 (772), PTPRT (1,026), and AGBL1 (1,068). For each gene of these nine, we resample the reference genotype matrix 5,000 times to increase the sample size, from the original 503 individuals, to yield more stable and realistic GWAS simulations.

#### Gene simulation

To simulate GWAS SNP-level statistics, we use a gene’s *n* × *m* reference genotype matrix **X** where *n* = 5,000 individuals, *m* = number of SNPs, and *X_i,j_* ∈ {0,1,2} is the number of effect alleles for individual *i* at SNP *j*. We generate simulations for null genes in the following way:

1. Compute the gene’s *m* × *m* correlation matrix **R**.
2. Determine the phenotype status *Y_i_* ∈ {0,1} for each individual *i* ∈ [*n*] depending on the expected case rate ***η***:

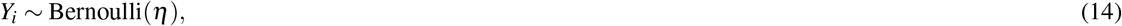
3. For each individual SNP *j* ∈ [*m*], fit a logistic regression model logit(P(*Y_i_* = 1|*X_j_*)) = *β*_0_ + *β_j_* · *X_j_* using all 5,000 individuals, returning the SNP’s two-sided p-value *p_j_*.
4. Compute gene-level p-values using the SNP-level statistics with each of the considered methods: (1) *MAGMA: paper*, (2) *MAGMA: ρ*^2^, (3) *MAGMA: code*, (4) *Two-sided approximation*, and (5) *Corrected*.

To measure the type 1 error rate at target *α* = 0.05 for the five different methods, we generate 1,000,000 independent null simulations for each of the nine example genes with the case-rate *η* = 0.5.

#### Multiple testing simulation

We assess each method’s multiple testing performance by simulating sets of null genes that are comparable in size to the number of genes tested in H-MAGMA. To make the simulations more realistic, we generate each set of genes based on the distribution of the number of SNPs assigned to each gene for the H-MAGMA autism spectrum disorder (ASD) results using the adult Hi-C data annotations. This corresponds to roughly 82.6% from [10,100], 16.4% (100,500], and 1.0% from (500,∞). Using the nine randomly picked genes from each of the size buckets, we construct each set of null genes by assigning:

- 14,663 from each gene with *m* ∈ [10,100],
- 2,827 from each gene with *m* ∈ (100,500],
- 177 from each gene with *m* ∈ (500, ∞).

We repeat this process to generate 1,000 sets of *G* = 53,001 null genes to assess each method’s impact on multiple testing corrections. First, we use the Bonferroni correction, rejecting the gene’s null hypothesis if its p-value is ≤ 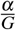, to control the target family-wise error rate (FWER), the probability of making at least one type 1 error, at *α* = 0.05. We then apply the Benjamini-Hochberg procedure^4^ (BH) to control the false discovery rate (FDR) at target *α* = 0.05, and compute the average false discovery proportion (FDP) across the simulations.

#### Gene-set analysis simulation

We measure the downstream impact of the different approaches for computing gene-level p-values on self-contained gene-set analysis, i.e., test whether a set of genes is associated with a phenotype. There are several ways for performing gene-set analysis^15^, but we consider three approaches for computing a test statistic *T_s_* for gene-set *s* with *G* genes:

1. MAGMA^1^ gene-set analysis: each gene *g*’s p-value *p_g_* is converted to a one-sided z-statistic, *z_g_* = *ϕ*^−1^(1 − *p_g_*), then the test statistic is

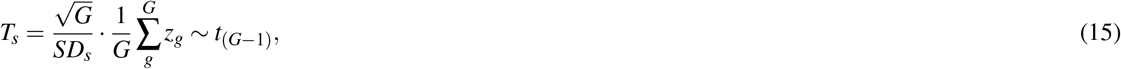

where *SD_s_* is the sample standard deviation of the gene-set’s one-sided z-statistics. The one-sided gene-set p-value is computed as *p_s_* = 1 − *F*_*t*_(*G*−1)__ (*T_s_*), where *F*_*t*_(*G*−1)__ is the cumulative distribution function for the *t*-distribution with *G* − 1 degrees of freedom. *Note: this is a separate test from the MAGMA gene-level test described earlier in this manuscript*.
2. Fisher’s combination test: as presented in Equation 1,

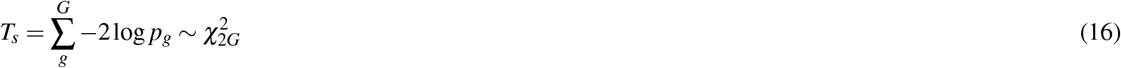
3. Stouffer’s z-test: similar to MAGMA, one-sided z-statistics, *z_g_* = *ϕ*^−1^ (1 − *p_g_*), are computed for the test statistic,

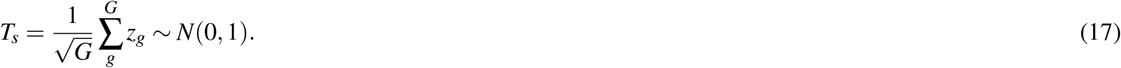 The gene-set p-value is then computed as *p_s_* = 1 − *ϕ*(*T_s_*).

Both Fisher’s and Stouffer’s are testing the global null of the gene-set,

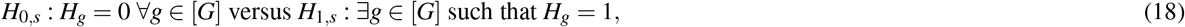

i.e. all of the individual genes, *g* ∈ [*G*] in gene-set *s*, are null versus at least one gene in the set is non-null. In comparison, the MAGMA approach is a one-sided test for whether the genes in gene-set s are jointly associated with the phenotype based on a measure of the set’s effect size *μ_s_*,

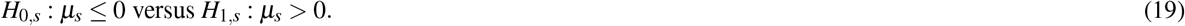

We simulate 200,000 gene-sets of size *G* = 45, constructed with five independent null genes from each example gene, to compare the gene-set analysis type 1 error rate that follows from using the different methods for computing gene-level p-values.

#### Replicating H-MAGMA Analysis

We assess the impact of correcting the usage of MAGMA for computing gene-level p-values with a Monte Carlo-based approach when applied to the SNP annotations used in H-MAGMA by Sey et al.^3^. Specifically, we recreate the results presented in Extended Data Fig. 1b of the H-MAGMA manuscript for the GWAS results of five psychiatric disorders: attention-deficit/hyperactivity disorder^16^ (ADHD), autism spectrum disorder^17^ (ASD), schizophrenia^18^ (SCZ), bipolar disorder^19^ (BD) and major depressive disorder^20^ (MDD). For each of the five GWAS results, Sey et al.^3^ assign SNPs to genes based on annotations derived from two different sources of Hi-C data: (1) adult and (2) fetal brain tissue samples (available on GitHub: https://github.com/thewonlab/H-MAGMA). H-MAGMA relies on MAGMA to compute the GWAS gene-level p-values for both adult and fetal Hi-C annotations, then proceeds to identify which genes are associated with the respective GWAS phenotype using BH to control FDR at target *α* = 0.05. The counts reported by Sey et al. in Extended Data Fig. 1b correspond to taking the union of the adult and fetal sets of BH discoveries.

We use the same set of Hi-C derived annotations to compute the *Corrected* Monte Carlo-based gene-level p-values with *N* = 1,200,000 null simulations for each gene. Each gene’s assumed covariance matrix is based on the sample of 503 individuals of European ancestry from the 1000 Genomes project, which is the same reference data used by Sey et al. in H-MAGMA. We proceed to identify the union of adult and fetal BH identified genes at target FDR level *α* = 0.05 for each of the five psychiatric disorders’ GWAS and compare to the reported H-MAGMA results. We encountered minor discrepancies in the total number of genes with p-values in a subset of the GWAS results as compared to H-MAGMA, since we were unable to compute gene-level p-values for the following: forty-one adult and forty-five fetal for ADHD, two adult and one fetal for ASD, and one fetal for SCZ. These differences are likely due to pre-processing steps that are not publicly available since our analysis matches the set of genes with p-values returned by using the MAGMA software directly, including the same set of SNPs by accounting for synonymous identifiers. However, these differences are negligible and do not impact our results as only one of these missing genes were reported as an association for ASD by H-MAGMA.

### Results

#### Comparison of type 1 error rate control

Fig. 1b and Supplementary Fig. 2 display the type 1 error rates at target *α* = 0.05, plus/minus two standard errors, for the example genes with each of the considered methods to compute gene-level p-values. The same pattern holds for each gene: the different variants of MAGMA display inflated type 1 error rates while the *Corrected* Monte Carlo-based approach maintains valid control. Unsurprisingly, the incorrect usage of Brown’s covariance approximation (*MAGMA: paper*) displays the greatest error rate inflation. *MAGMA: code* yields comparable results to the *Two-sided approximation*, however, it appears to display higher inflated error rates with more severity for larger genes.

These results are consistent with recent analysis regarding the behavior of Brown’s approximation when applied to two-sided tests generated from multivariate normal data^14^, whereas our simulations use real genotype data rather than rely on distributional assumptions. The reason for this poor performance stems from the failure of Brown’s re-scaled *χ*^2^ distribution to properly fit the actual null test statistic distribution, regardless of the covariance approximation, as seen in Fig. 1a for the ‘toy’ multivariate Gaussian example with *m* = 50.

#### Impact on multiple testing

Supplementary Fig. 3 displays the observed FWER across 1,000 simulations of applying the Bonferroni correction to sets of 53,001 null genes, by each gene-level method. Only the *Corrected* Monte Carlo-based approach controls FWER at the target rate *α* = 0.05 (indicated by the dashed red line), while all of the other MAGMA variants do not maintain valid control within plus/minus two standard errors. The *MAGMA: code* approach appears to be more conservative than using the *Two sided approximation* approach, likely from its adjustment power *c*. For context, Supplementary Fig. 4a displays the proportion of simulations with zero to six false positives with the Bonferroni correction for each method (excluding *MAGMA: paper*) while Supplementary Fig. 4b displays the distribution of the number of false positives represented by boxplots, with points denoting single simulation results, for each method (dashed red line indicates one observed false positive). The *Corrected* Monte Carlo-based approach is the only method that yield less than one false positive in the vast majority of simulations (as implied by the FWER control), while the other methods display a substantial number of simulations resulting in one to six false positives or, in the case of using the *MAGMA: paper* approach, between two to three hundred false positives. We observe similar patterns of inflation when applying BH for FDR control as displayed in Fig. 1d.

We investigate this further by examining the p-value distributions for the simulated sets of 53,001 genes. Fig. 1c displays histograms for the null gene-level p-values by method averaged over 1,000 simulations (standard errors are too small to be visible), with the black horizontal line denoting the ideal uniform distribution for null p-values. However, only the *Corrected* null p-value distribution displays such behavior. The different MAGMA variants, including the *Two-sided approximation*, display an inflation in smaller p-values along with non-uniform p-value distributions, such as fewer than expected number of genes for the bin of largest p-values (≥ 0.95). The observed inflated FWER and FDR in Supplementary Fig. 3 and Fig. 1d emphasize the failings of using these improper, non-uniform null distribution in typical multiple testing procedures.

#### Impact on gene-set analysis type 1 error control

The type 1 error rate for the gene-set simulations are displayed in Supplementary Fig. 5. Regardless of the gene-set analysis method, only the *Corrected* Monte Carlo-based approach displays control at the at the target type 1 error rate *α* = 0.05 (as indicated by the dashed red line). The behavior for the other methods varies depending on the type of gene-set analysis. The different types of MAGMA gene-level p-values result in overly conservative results for the MAGMA gene-set analysis approach, while they display inflated error rates for both Fisher and Stouffer’s method to varying degrees (with the notable exception that *MAGMA: code* is conservative with Stouffer’s method).

The difference in behavior of the gene-set error rate control is driven by the non-uniform p-value distribution. Supplementary Fig. 6 displays a comparison of the resulting gene-set null p-value distributions by gene-set analysis method (columns) and method for computing gene-level p-values (rows). We clearly see that for both the MAGMA and Stouffer’s gene-set analysis methods, the *MAGMA: code* approach yields conservative p-value distributions. To further investigate this behavior, Supplementary Fig. 7 displays the resulting MAGMA gene-set analysis *z_s_* distributions by the gene-level method, with the expected *t*_(*G*−1)_ (where *G* = 45) distribution curve overlaid in red. We see that both the *MAGMA: paper* and *MAGMA: code* approaches are shifted to the left with more negative values. The use of the adjustment power *c* in the *MAGMA: code* approach, which applies a more conservative adjustment as the p-value decreases, has a noticeable downstream effect on the MAGMA gene-set analysis results. The Monte Carlo-based approaches are the only gene-level methods yielding appropriate null distributions and results regardless of the gene-set analysis method.

#### Results for H-MAGMA Replication

Supplementary Table 1 and Fig. 2b display the substantial reduction in the number of discoveries reported by H-MAGMA after using the *Corrected* Monte Carlo-based approach. Fig. 2a highlights the improper H-MAGMA ASD p-value distribution (based on adult Hi-C data annotations) consistent with behavior observed in our simulations (Fig. 1c).

**Supplementary Fig. 2.**
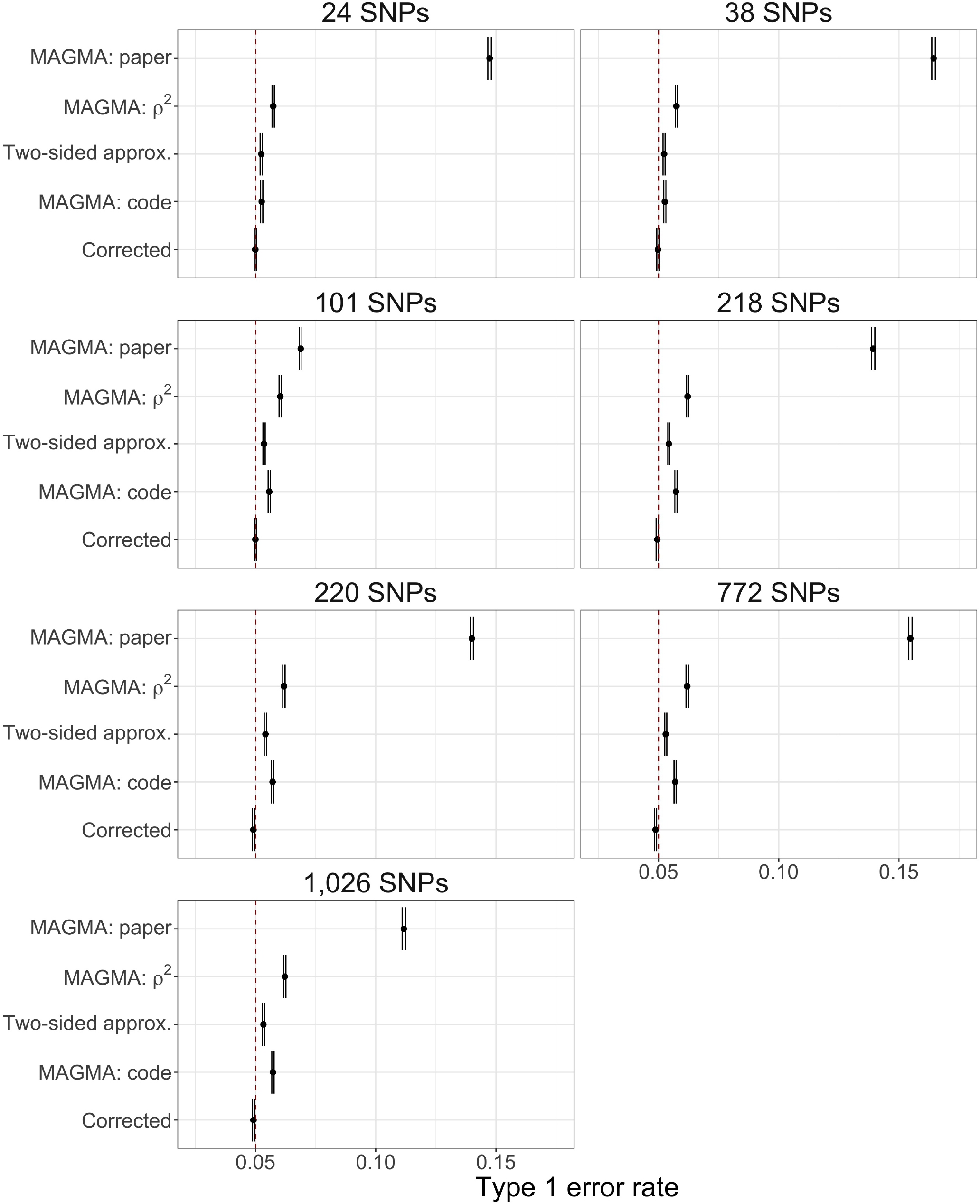
Comparison of the type 1 error rate control, plus/minus two standard errors, at target *α* = 0.05 (denoted by dashed red line) by gene-level method for seven example genes of different sizes.

**Supplementary Fig. 3.**
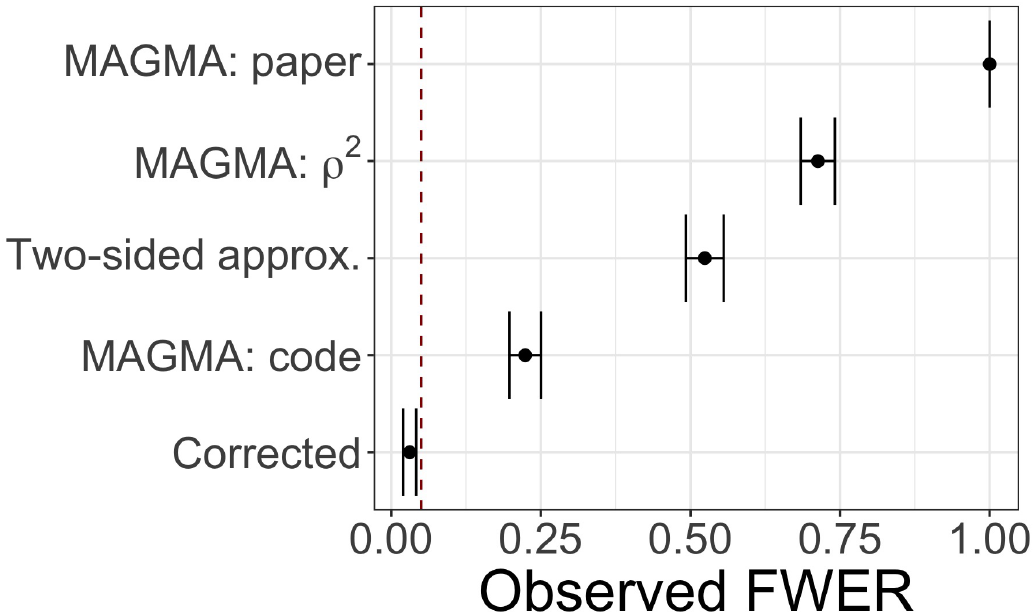
Comparison of multiple testing family-wise error rate (FWER), plus/minus two standard errors, at target *α* = 0.05 (denoted with dashed red line) by gene-level method.

**Supplementary Fig. 4.**
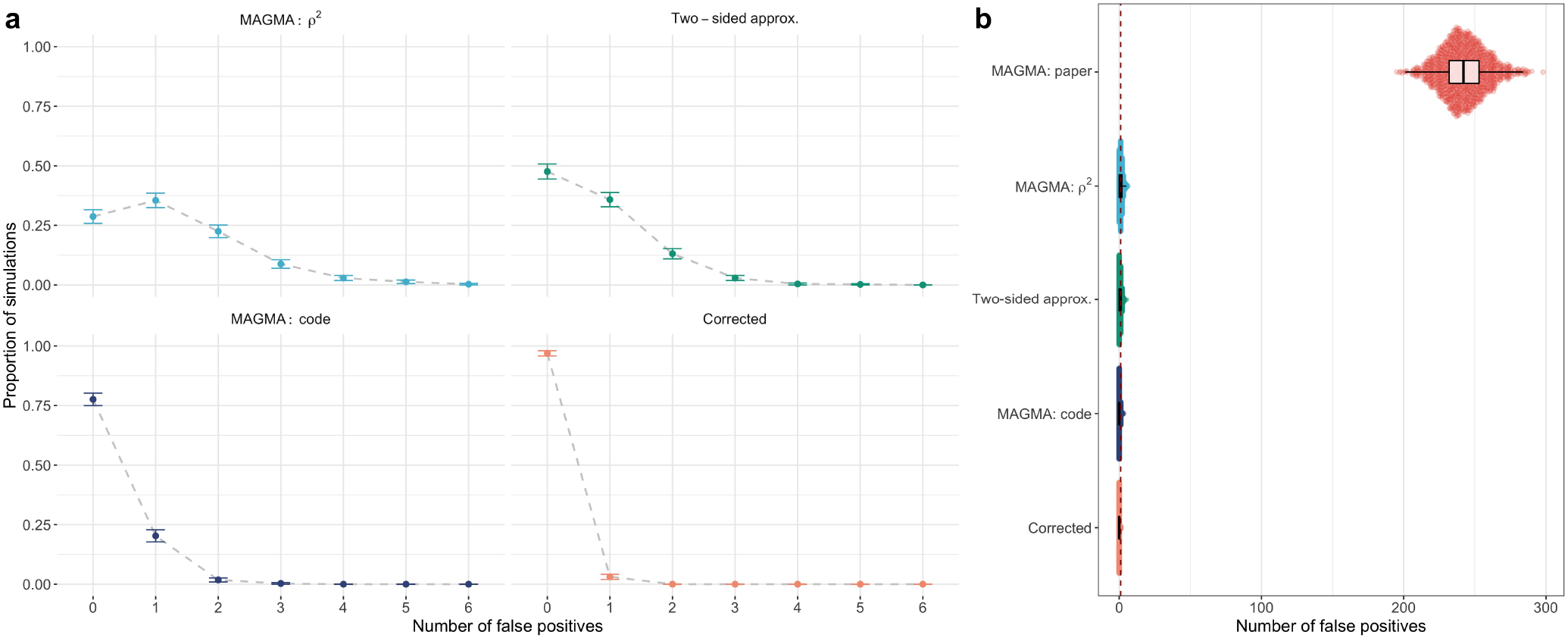
**a**, Comparison of the proportion of simulations with zero to six false positives for each method, excluding *MAGMA: paper*, using the Bonferroni correction at target *α* = 0.05 (plus/minus two standard errors). **b**, Distribution of the number of false positives represented by boxplots, with points denoting single simulation results, for each method (dashed red line indicates one observed false positive).

**Supplementary Fig. 5.**
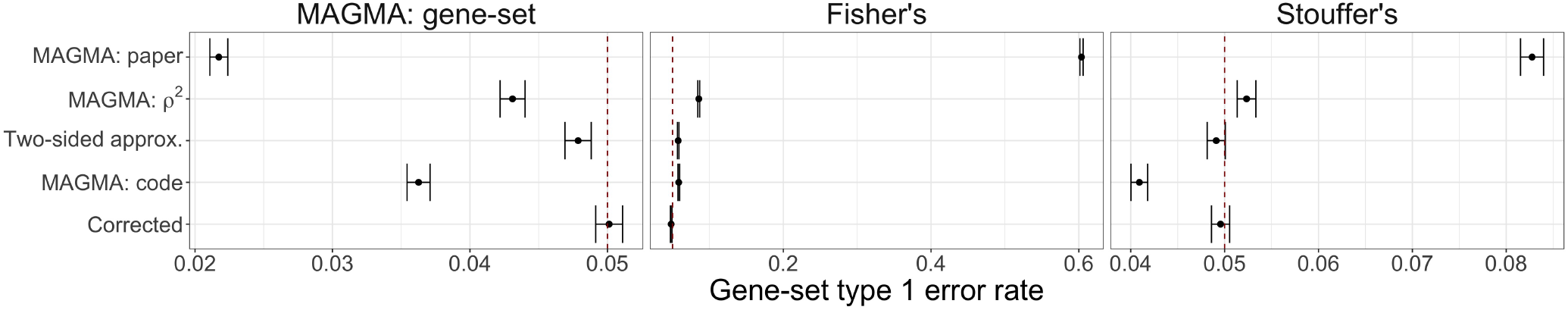
Comparison of gene-set analysis type 1 error, plus/minus two-standard errors, at target *α* = 0.05 (denoted with dashed red line) by gene-level method for each considered gene-set analysis method (from left to right) MAGMA: gene-set, Fisher’s combination test, and Stouffer’s z-test.

**Supplementary Fig. 6.**
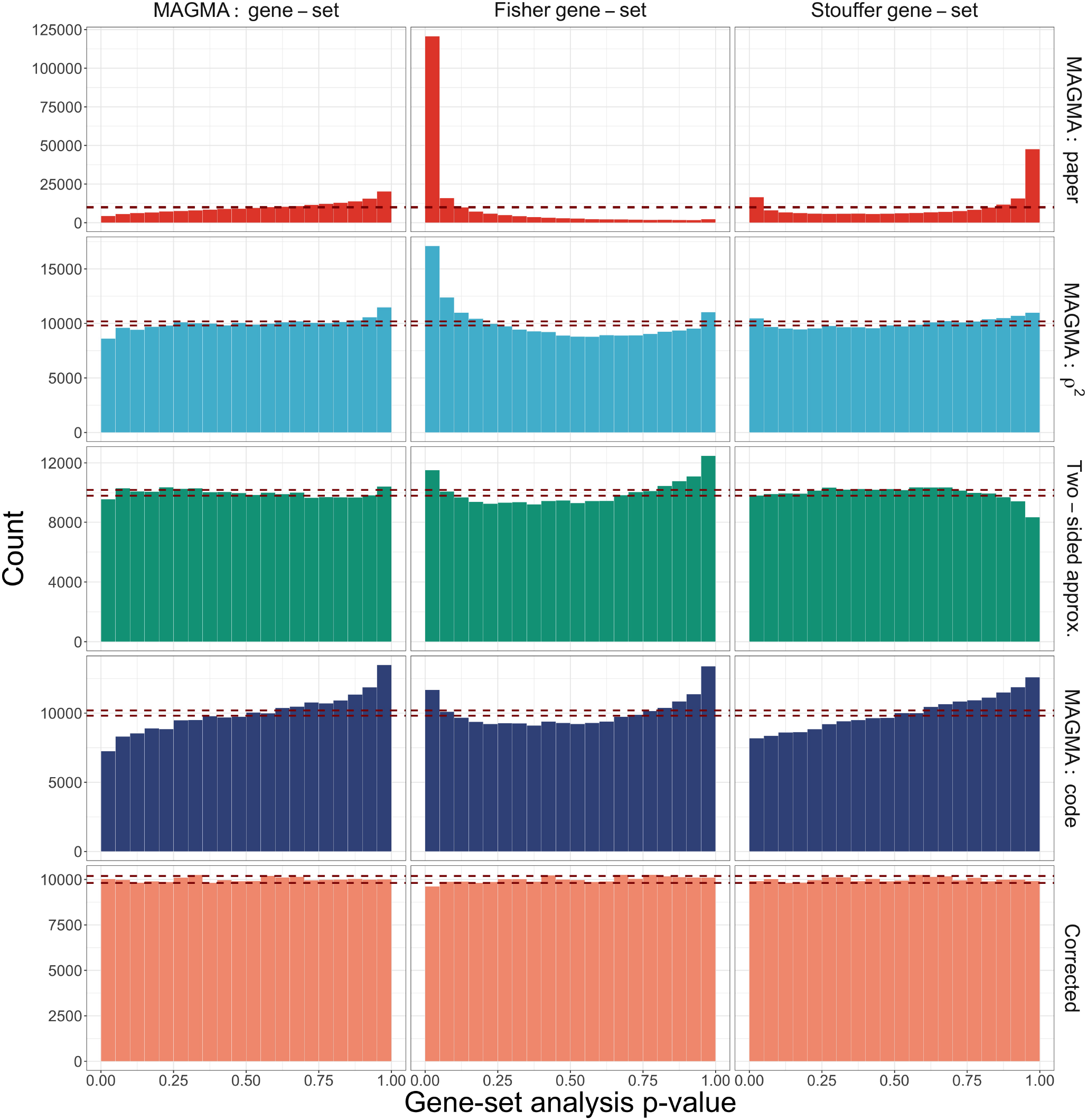
Comparison of null histograms by gene-set analysis method (columns) and methods for computing gene-level p-values (rows) for gene-sets of size *G* = 45 genes. Red dashed lines indicate expected 2.5% and 97.5% quantiles for uniform p-values across 200,000 simulations.

**Supplementary Fig. 7.**
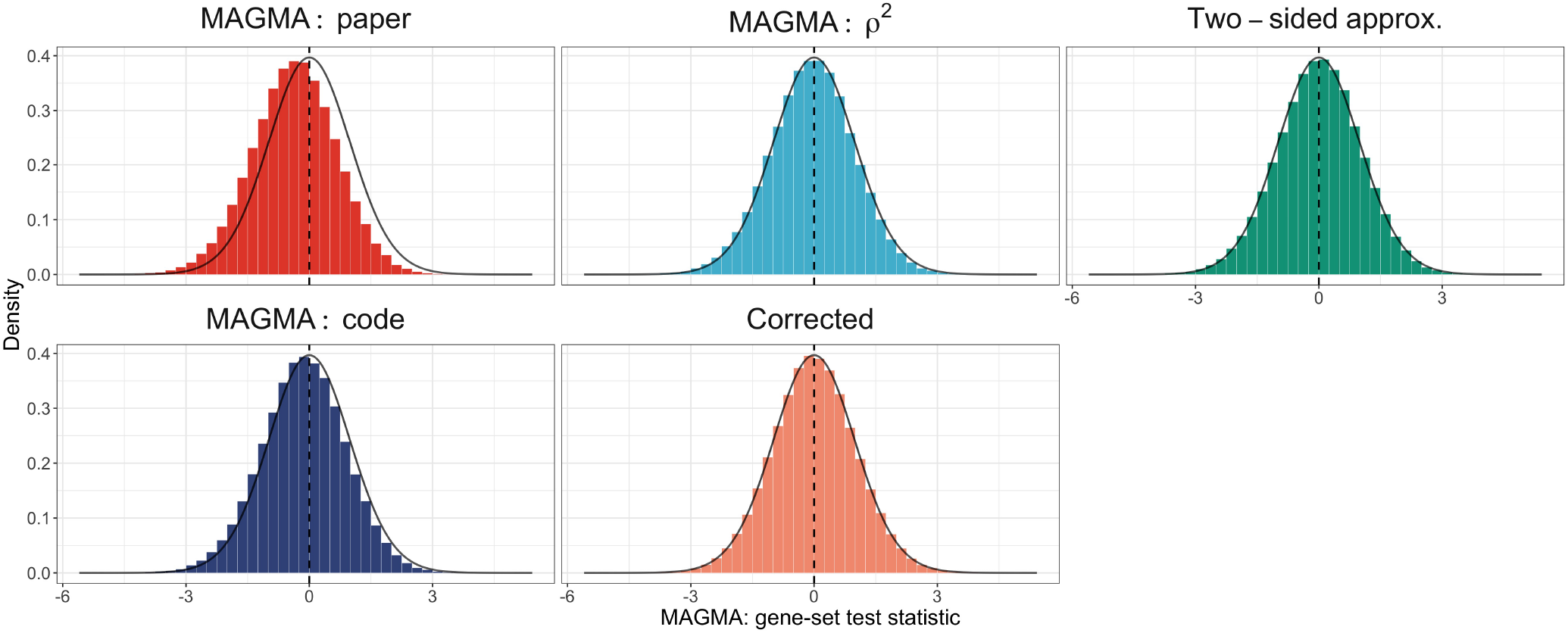
Comparison of distributions for MAGMA gene-set analysis standardized test statistic by methods for computing gene-level p-values. Red density curve displays the expected *t*_(*G*−1)_ distribution, with the red vertical dashed line denoting the center at zero.

**Supplementary Table 1.** Comparison of number of BH discoveries (union across adult and fetal Hi-C data annotations) with H-MAGMA versus corrected Monte Carlo-based approach.

| Phenotype | H-MAGMA | Corrected |
| --- | --- | --- |
| ADHD | 486 | 223 |
| ASD | 275 | 125 |
| BD | 1,931 | 1,111 |
| SCZ | 9,217 | 8,066 |
| MDD | 3,167 | 2,201 |

## Acknowledgements

This work was supported by National Institute of Mental Health Grant R37MH057881, the Simons Foundation Grant SFARI SF575097, and the National Science Foundation DMS 1613202 Grant.

## Author contributions statement

R.Y., M.G., K.R., and B.D. conceived the experiments; R.Y. conducted the experiments; and R.Y., M.G., K.R., and B.D. wrote the manuscript.

## Additional information

### Accession codes

None.

### Competing interests

The authors declare no competing interests.

## Notes

https://github.com/ryurko/HMAGMA-comment

### Competing Interest Statement

The authors have declared no competing interest.

